# Identification of a novel papillary renal cell carcinoma molecular subtype characterized by HIF-pathway over-activation

**DOI:** 10.1101/2025.01.24.632989

**Authors:** Javier de Nicolás-Hernández, Alicia Arenas, Carlos Valdivia, María Santos, Javier Lanillos, Nicole Bechmann, Mirko Peitzsch, Eduardo Gil-Vilariño, María Monteagudo, Sara Mellid, Cristina Montero-Conde, Alberto Cascón, Luis Javier Leandro-García, Guillermo de Velasco, Ignacio Durán, Javier Puente, Jesús García-Donas, Eduardo Caleiras, Mercedes Robledo, Cristina Rodríguez-Antona

## Abstract

**Purpose:** Papillary renal cell carcinoma (pRCC), the second most common subtype of renal cancer, exhibits heterogeneity in molecular features and response to targeted therapies, including antiangiogenic drugs. Discovering molecular biomarkers able to stratify pRCC patients into clinically relevant subgroups and understanding the underlying mechanisms are urgently needed to advance precision medicine. Here, we molecularly dissect a large pRCC series through the expression of the targets of hypoxia inducible factors (HIF), master regulators of angiogenesis, metabolic reprogramming and immune microenvironment.

**Experimental Design:** We merged and analyzed multi-omic and clinical data of 346 patients derived from two pRCC series (TCGA-KIRP and a Spanish metastatic series). Altered pathways and differences in tumor microenvironment were identified through tumor transcriptomic analyses. Tumor metabolome analysis was performed in selected cases.

**Results:** Molecular revision of driver mutations classified 302 patients as pRCC, while uncovering misclassified cases. Analysis of HIF targets gene expression identified a subset of pRCC tumors with increased HIF activity (31%; “HIF-active”). These tumors were characterized by high hypoxia scores, increased angiogenesis, low expression of Krebs cycle genes and mitochondrial activity, high immune infiltration and increased epithelial-mesenchymal transition (P<0.0001; each feature) and they were associated with increased metastasis risk and worse overall survival (P<0.005). Metabolome analyses revealed that HIF-active tumors accumulated L-2-hydroxyglutarate (L-2-HG), an oncometabolite that leads to pseudohypoxia by inhibiting HIF prolyl hydroxylases and preventing HIFα degradation. L-2-HG-accumulation agreed with diminished *L2HGDH* expression, associated with losing one copy of the gene and low expression of *PPARGC1A*, which regulates its transcription.

**Conclusions:** HIF-pathway overactivation defines a pseudohypoxic pRCC molecular subgroup characterized by high angiogenesis, immune infiltration and aggressive features. These findings advance our understanding of pRCC diversity, revealing potential clues for the variable response of patients to targeted therapies and paving the way to more personalized pRCC treatments.

## INTRODUCTION

Papillary renal cell carcinoma (pRCC) is the second most common type of renal cell carcinoma (RCC) accounting for 10-15% of diagnoses^1^ Traditionally, pRCC followed a dichotomic type 1 or type 2 classification^1,2^; however, in 2022 the World Health Organization abandoned this simplistic division and recognized that the spectrum of pRCC is more complex^3^. In this regard, the differential diagnosis of pRCC now excludes several new emerging entities with specific clinical and molecular characteristics including FH-deficient, SDH-deficient, ALK-rearranged, TFE3-rearranged and TFEB-altered RCC, some of which can exhibit papillary morphology^3^. Despite these advances, there is still significant molecular and clinical heterogeneity within pRCCs that remains unresolved^1,2^.

While surgical resection in patients with localized pRCC may be curative, metastatic disease presents a major clinical challenge. Due to the relatively low incidence of pRCC and its molecular diversity, therapeutic options are mainly unselective, as they rely on clinical trials performed with metastatic clear cell RCC (ccRCC), the most common histologic subtype. Unlike pRCC, ccRCC is homogenously driven by mutations in the von Hippel-Lindau (VHL) tumor suppressor, which causes abnormal hypoxia-inducible factor (HIF) accumulation^4^. Current frontline treatments for metastatic RCC of any histology include anti-angiogenic tyrosine kinase inhibitors (TKIs) and immune checkpoint inhibitors (ICIs)^5^. However, response rates of metastatic pRCC patients to these drugs are heterogeneous and frequently suboptimal^6–8^, highlighting the diversity of papillary tumors. Targeted therapies specific to pRCC have shown unclear results so far, exemplified by inhibitors of MET⁹, a controversial target found in less than 15% of pRCC cases¹. Therefore, there is still an urgent need to identify new biomarkers able to stratify pRCC patients according to molecular alterations that can guide treatment selection.

The HIF family of transcription factors are the key regulators of cellular adaptation to low oxygen^10,11^. The degradation of the HIFα subunit in the presence of oxygen is mediated by the coordinated action of HIF prolyl-hydroxylases (PHDs)^12^ and the E3-ubiquitin ligase VHL^13^. Some tumors exhibit a phenomenon called pseudohypoxia, in which HIFα accumulates independently of oxygen levels, and promotes tumorogenesis^14^. The most common example of pseudohypoxia in RCC occurs in ccRCC through *VHL* mutation^4^. The pivotal role of HIF in ccRCC explains the sensitivity of this tumor to anti-angiogenic drugs and, more recently, to the new HIF2α inhibitors. Pseudohypoxia also occurs in the rare fumarate hydratase (FH) and succinate dehydrogenase (SDH) deficient RCC entities. In these cases, the tumor-specific loss of FH or SDH activity leads to the accumulation of fumarate and succinate oncometabolites, respectively, which strongly inhibit HIF PHDs^15^.

pRCC lacks driver mutations in *VHL*, *FH* or *SDH*, suggesting low HIF pathway activation. However, Lombardi et al. developed a pan-cancer method to infer HIF activation, and a significant variability among pRCCs can be observed in this data, with some tumors showing high “hypoxia scores” ^16^. This variable HIF activation is unexplored and may be associated with the molecular heterogeneity of papillary tumors. Moreover, this variability in HIF activation could also explain why only a fraction of pRCC patients benefit from anti-angiogenic drugs, with objective response rates ranging from 9 to 30% ^1,6–9^.

In this study, using a multi-omics approach in a large pRCC cohort, we define a tumor subgroup with high HIF activity and conduct its molecular and clinical characterization, aiming at providing novel tools to perform biologically-driven drug selection for metastatic pRCC patients. Furthermore, we also elucidate the mechanisms underlying HIF overactivation.

## MATERIALS & METHODS

### Sample series

Spanish series sample inclusion required papillary histology annotation in the anatomopathological report or the clinical record of the patient. The study included a total of 66 formalin-fixed paraffin-embedded (FFPE) tumor samples from 55 metastatic patients, collected by 16 hospitals across Spain. Written informed consent was obtained from all patients, and the study was approved by the corresponding ethical review boards and complied with the declaration of Helsinki.

The Cancer Genome Atlas kidney papillary cell carcinoma (TCGA-KIRP) cohort was composed of 292 primary tumors, corresponding to 291 patients, from which the genomic data and clinical phenotype was downloaded from The Cancer Genome Atlas (TCGA) portal (https://gdc.cancer.gov)^2^. Overall survival (OS) data was extracted from cBioPortal^17^ (TCGA-KIRP Firehouse Legacy database).

The clinical and molecular characteristics of the combined series are provided in Supplementary Table 1.

### Nucleic acid extraction and quality control

Tumor samples from the Spanish series were reviewed by a pathologist (E. Caleiras). DNA and RNA were isolated from samples with >75% cancer cells. Tumor DNA was obtained with truXTRAC FFPE DNA (Covaris, 520136) and tumor RNA using Maxwell RSC RNA FFPE Kit (Promega, AS1440). Germline DNA was extracted either from peripheral blood mononuclear cells using Maxwell RSC Whole Blood DNA Kit (Promega, AS1520) or from tumor-adjacent normal tissue using truXTRAC FFPE DNA. DNA and RNA was quantified using Quantus Fluorometer (Promega) and integrity was examined using a High Sensitivity DNA Kit (Agilent, 5067-4626) or an RNA 6000 Nano Kit (Agilent, 5067-1511) in a 2100 Bioanalyzer Instrument (Agilent).

### DNA sequencing

In the Spanish series, germline DNA library preparation was performed as previously described^18^. The tumor DNA was sequenced using custom panels designed to target the coding region of 33 genes implicated in RCC and cancer. These genes are detailed in Supplementary Table 1. In brief, for library preparation, SeqCap EZ Choice Enrichment Kit (Roche) or Twist Library Preparation EF Kit 1.0 (Twist) were used according to the manufacturer’s instructions using 250–500 ng of DNA. Sequencing was performed in a HiSeq or NextSeq sequencer (Illumina) configured to generate 100 bp paired-end reads. A total of 65 tumor samples were successfully sequenced with a median bait coverage of 304 (min–max: 53-823; interquartile range [IQR]: 193-382). For alignment, GRCh37/hg19 assembly was used as reference using BWA_mem2 v2.2.1 tool^19^ and for the calling of somatic variants Mutect2 v2.2^20^ (GATK4 v4.2.5.0) was used. Ensembl Variant Effect Predictor annotation tool (VEP) v100 was used to predict variant impact and only variants with “high” and “moderate” impact were considered for the analysis. Somatic variants with a MAF > 0.00001 in gnomAD were filtered out. Variants with <10 total reads, <4 altered reads or a fraction of altered reads <0.15 were excluded. For low-quality sequencing samples, the VAF threshold was set up to 0.25. Additionally, ABRA v0.97 software^21^ was employed to identify *TFE3*, *TFEB* and *ALK* translocations. Low confidence variants were reviewed using the Integrative Genomics Viewer (IGV) and manually curated. For copy number alteration (CNA) detection, FACETS v0.5.14 algorithm^22^, using R package *cnv-facets* v0.15.0, was employed and the 14q chromosome status of the tumors from the Spanish collection was inferred.

For TCGA-KIRP, germline BAM files were downloaded from the TCGA repository at NCI Genomic Data Commons (https://portal.gdc.cancer.gov/) (dbGaP accession ID phs000178). The GRCh38 genome assembly was then used as a reference for the alignment. Variant calling was performed using Haplotypecaller^23^, followed by VEP annotation. Somatic alterations of TCGA-KIRP were extracted from cBioPortal Pancancer Database. For CNA analysis, Gistic2 Gene level-copy number values of TCGA-KIRP cohort were downloaded from UCSC Xena Browser (https://xenabrowser.net/)^24^.

### RCC subtype molecular revision

For RCC subtype molecular revision, the following genes were analyzed: i) *VHL*, *FH*, *SDHA/B/C/D* for germline variants, or ii) *VHL*, *FH*, *SDHA/B/C/D, TFE3, TFEB, ALK, SMARCB1, ELOC* for somatic variants. Alterations categorized as driver pathogenic variants in genes associated with RCC entities different from pRCC led to subtype reclassification (see Suppl. Tables 1 and 2).

In the Spanish series, germline pathogenic variants in *VHL*, *FH* and *SDHA/B/C/D* were detected as previously described^55^. Somatic alterations required VAF >0.35 to be considered drivers. In the TCGA-KIRP series, germline variants were considered driver mutations when they were: i) classified by ClinVar as pathogenic or likely pathogenic, or ii) they were rare loss-of-function variants (gnomAD MAF<0.00001). In addition, *FH* rare missense or inframe variants in the presence of CpG island methylator phenotype (CIMP) were also classified as driver mutations. For somatic drivers, we included mutations categorized as oncogenic by OncoKB^TM^, *TFE3* fusions, *TFEB* fusions or amplifications, *ALK* fusions, and gene homodeletions with mean CNA log2 value <-2.

### DNA Methylation data

Methylation450K beta values for the available TCGA-KIRP cases were downloaded from UCSC Xena Browser (https://xenabrowser.net/)^24^. Probes corresponding to SNVs were removed, as well as probes with NA value in any sample (n=392,845 remaining probes). We then kept probes that were unmethylated (mean β-value <0.1) in normal tissue (n=110,553 remaining probes) and, finally, we chose probes with a standard deviation greater than 0.1 (n=14,915 final probes).

### RNA sequencing

In the Spanish series, cDNA libraries from FFPE tumors were prepared using QuantSeq 3’ mRNA-Seq Library Prep Kit FWD for Illumina (Lexogen, 015), following the vendor’s protocol for low input/ low quality/ FFPE RNA. 200–500 ng of total RNA was used as starting material and the PCR Add-on Kit for Illumina (Lexogen, 020) was used to adjust the number of cycles for library amplification. Libraries were sequenced on a NovaSeq6000 (Illumina). Image analysis, per-cycle base calling and quality score assignment was performed with Illumina Real Time Analysis software. Demultiplexing of BCL files to FASTQ format was performed with the bcl2fastq Software (Illumina). Expression counts were generated with BlueBee® Genomics Platform (Lexogen, 090-094), from FASTQ files obtained from QuantSeq platforms. STAR v2.5.2a^25^ algorithm was used to map the reads to the reference genome GRCh37. Samples failing quality control were excluded: samples with fewer than 500,000 reads and at least one of the following criteria: >45.000 genes with zero reads or a ratio “Reads not assigned to annotated genes/Gene reads” > 2.

For the TCGA-KIRP cohort, RNA-Seq HTSeq-Counts were downloaded from UCSC Xena Browser (https://xenabrowser.net/)^24^.

### RNA-seq data normalization, clustering, score calculations and gene/pathway analyses

Gene expression counts from the Spanish and TCGA-KIRP series were integrated by common gene symbols using ComBat-seq^26^ from R/Bioconductor package *sva* v.3.46.0^27^ on log-transformed values and normalization was performed using DESEQ2 v1.38.3^28^.

For HIF-target genes clustering, we applied optimal K-means clustering (K=2) based on the expression of 461 HIF-target genes (GSEA_Hallmark_Hypoxia_M5891; GSEA_Angiogenesis_M14493; Selected hypoxia-inducible transcription factor target genes in ccRCC^29^, hypoxic cluster genes in pheochromocytoma^30,31^; Supplementary table 3) using R *NbClust*^32^ and *cluster*^33^ packages.

The KIRP-adjusted Hypoxia Score was calculated using the KIRP-best correlating genes provided in Lombardi et al.^16^. In short, we quantile normalized the data for each gene (i.e. normalized the data so that the average expression for each gene was 0 and the standard deviation was 1) to avoid the score being dominated by the most highly expressed or variable genes. We also calculated the Hypoxia Score based on the defined pan-cancer 48-gene signature^16^.

The epithelial-mesenchymal transition (EMT) score was calculated as previously described in Aggarwal RK et al.^34^. The combined TCGA-KIRP plus Spanish datasets were analyzed for these genes and based on the median value of the composite EMT score, samples classified as “high” and “low” EMT.

Differential gene expression analysis between groups defined by HIF activity was performed using *limma* tool within the Phantasus v1.21.5 web tool^35^. GSEA pre-ranked analysis was executed with GSEA software v4.2.3^36^ (RRID:SCR_003199), using the ‘H: hallmark gene sets’ and ‘C5: ontology gene sets’ collections from the Molecular Signature Database (MSigDB; https://www.gsea-msigdb.org/gsea/msigdb/). Default settings were used except for collapse set to No_Collapse, Enrichment_statistic set to weighted, Max_size set to 500, Min_size set to 15 and Plot_Top_x set to 100. Input files were obtained from the differential expression analysis and ranked according to the *t* statistics.

### Tumor microenvironment (TME) analyses based on RNA-seq data

To calculate tumor purity, normalized counts were used to infer tumor purity for each sample using the estimate score provided by ESTIMATE algorithm v1.0.1336^37^. For TME subtype identification, gene set variation analysis (GSVA) enrichment scores for the 29 curated gene signatures (Fges) from Bagaev et al.^38^ were calculated using the GSVA R package^39^ on the COMBAT-seq normalized expression matrix. Tumors were then classified into two separate TME, named “Angiogenic” or “Non-Angiogenic” applying unsupervised k-means clustering (Euclidean distance, 1000 iterations) using Morpheus (https://software.broadinstitute.org/morpheus), similarly to the initial publication^38^.

Absolute abundance of 22 different immune cell subtypes was calculated using CIBERSORTx^40^ in each sample, using the COMBAT-seq normalized expression matrix. CIBERSORTx analysis parameters were the following: i) gene signature matrix: LM22, containing gene expression profiles of 22 immune cell subsets; ii) 1000 permutations were performed to obtain robust and reliable results; iii) B-mode batch correction to account for any potential batch effects and iv) quantile normalization was disabled during the analysis.

### Measurement of tissue Krebs cycle metabolites

Krebs cycle and related metabolites of formalin-fixed paraffin-embedded (FFPE) tumor tissue specimens were analyzed by ultrahigh pressure liquid chromatography with tandem mass spectrometry (UPHPLC-MS/MS) as previously described^41^. Metabolite ratios allow comparisons between samples.

### Statistical Analyses

All statistical analyses were performed using GraphPad v10.0.0 for Windows, (GraphPad Software, Boston, Massachusetts USA, www.graphpad.com). Differences were considered statistically significant if the p value was less than 0.05.

## RESULTS

### RCC molecular revision and histology reclassification

To ensure alignment with the WHO 2022 classification, germline and somatic alterations present in the Spanish and TCGA-KIRP series were analyzed, and tumors with driver mutations in genes defining RCC subtypes other than pRCC were excluded from the study. In TCGA-KIRP, this procedure revealed 10 FH-deficient RCC with mutations in *FH*, from which 5 cases had previously gone undetected (Supplementary Table 2). Additionally, we uncovered 3 primary tumors with driver mutations in *VHL* (indicating ccRCCs), 8 *TFE3-*translocated RCCs, 4 *TFEB*-rearranged RCCs (one of which is among the FH-deficient), 5 *SMARCB1*-deficient RCCs, 2 *ALK*-translocated RCCs and 1 *ELOC*-mutated RCC (Figure S1A). In the Spanish series, 4 *FH*-deficient, 2 *SDHB*-deficient, and 1 *TFE3*-translocated RCC cases were found to be originally misclassified as pRCC (Figure S1B).

After RCC subtype reclassification, the final pRCC series analyzed in this study comprised 302 patients and 303 independent primary tumors with good-quality transcriptomic data available (Figure S2). Demographic and clinical characteristics of remaining cases are provided in Table 1.

**Table 1.** Clinical characteristics of the pRCC patients included in the study.

| **Characteristics** | **Spanish series**<br>**(n=45) Nr (%)** | **TCGA-KIRP**<br>**(n=257) Nr (%)** |
| --- | --- | --- |
| **Age at diagnosis (years)** |  |  |
| Median [min, max] | 64 [31, 86] | 62 [28, 88] |
| **Sex** |  |  |
| Female | 12 (27) | 62 (24) |
| Male | 30 (67) | 195 (76) |
| Not Available | 3 (7) | - |
| **Stage at diagnosis** |  |  |
| Stage I | 6 (13) | 181 (70) |
| Stage II | 4 (9) | 25 (10) |
| Stage III | 8 (18) | 39 (15) |
| Stage IV | 20 (44) | 10 (4) |
| Not Available | 7 (16) | 2 (1) |
| **Developed Metastasis** |  |  |
| Yes | 45 (100) | 10 (4) |
| No | - | 188 (73) |
| Not Available | - | 59 (23) |
| **Vital Status** |  |  |
| Alive | 13 (29) | 222 (86) |
| Dead | 24 (56) | 35 (14) |
| Not Available | 7 (16) | - |
| **Overall survival time (months)** |  |  |
| 75% percentil [typical error] | 27.3 [7.5] | 76.6 [16.6] |
| 50% percentil [typical error] | 50.0 [15.4] | Not reached |

Pseudohypoxic RCC tumors identified by *FH* and *SDHB* pathogenic variants were used as a control group in the study (n=16).

### HIF transcriptional activity in pRCC

After combining the RNA-seq data of Spanish and TCGA-KIRP series with COMBAT-seq and ensuring a uniform distribution of samples (Fig. S3A-B), we employed a list of HIF targets to identify pRCC tumors with elevated HIF transcriptional activity. Using the Silhouette method, we determined that the optimal number of K-means clusters was 2 (K=2) and subsequently divided the samples into two groups (Fig. S4A-B). The pRCC cluster that included nearly all FH– and SDHB-deficient control tumors (n=16) was designated “HIF-active” (n=94) while the other was designated “Cluster A” (n=209) (Fig. 1A). Accordingly, tumors the HIF-active cluster exhibited higher hypoxia scores and were characterized by significantly increased expression levels of *HIF1A*, *EPAS1* and *HIF3A* (encoding for HIF1α, HIF2α and HIF3α, respectively, when compared to Cluster A (Fig. 1B-C, S4C-D). These features resemble those of the pseudohypoxic FH/SDH tumors.

**Figure 1.**
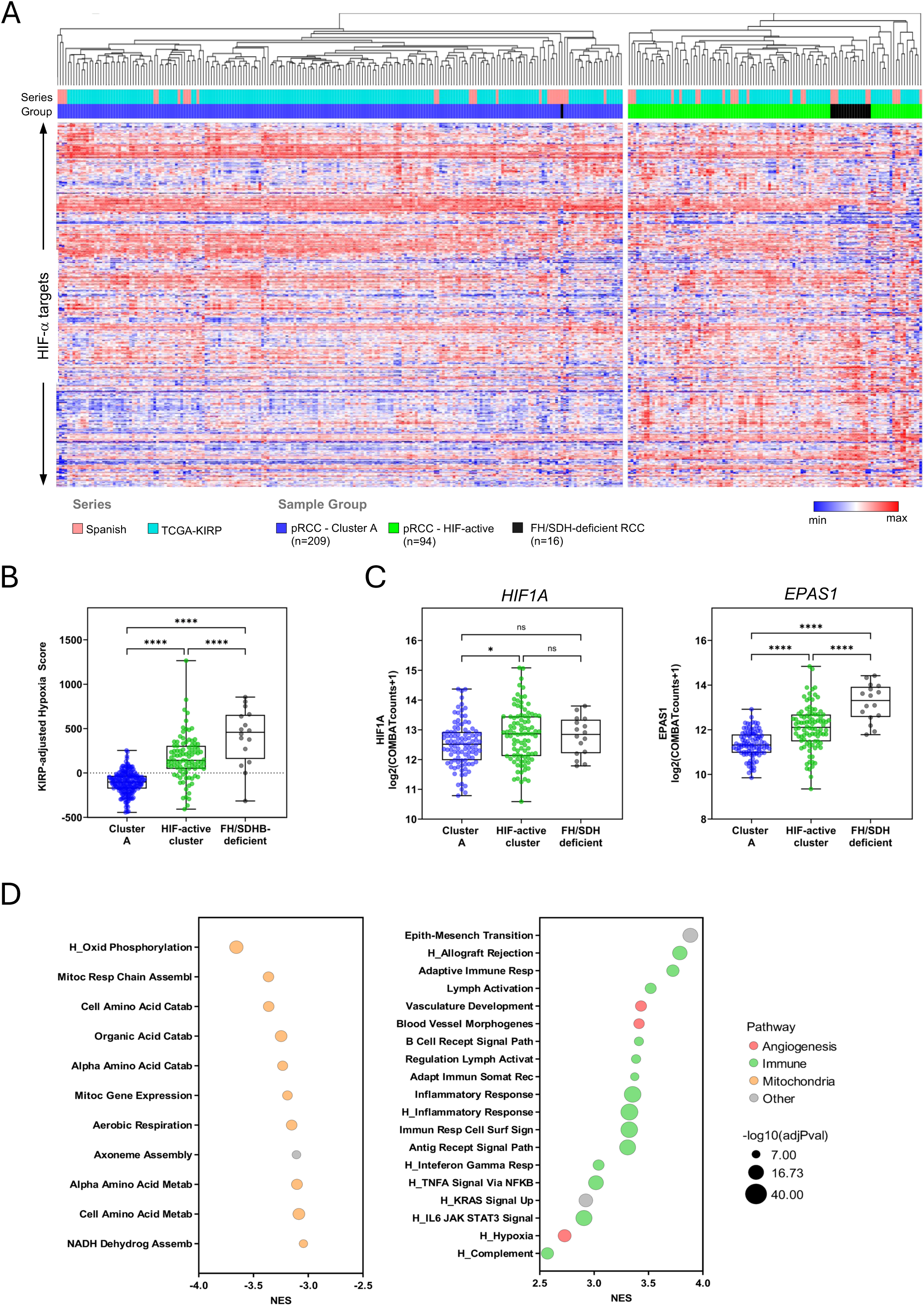
Transcriptomic characterization of HIF-active pRCC tumors. **A)** Heatmap representation of the clustering of 461 HIF-target genes. Tumors were annotated for sample series (Spanish, TCGA-KIRP) and sample group (pRCC-Cluster A, pRCC HIF-active, *FH/SDHB* deficient RCC). **B)** Hypoxia scores adjusted for genes positively correlating with HIF-metagene in TCGA-KIRP, as described by Lombardi *et al*. 2022. **C)** mRNA levels of *HIF1A* (left panel) and *EPAS1* (right panel) genes. Tukey’s multiple comparisons test. **D)** Bubble plot showing the top 10 down-regulated (left panel) and top 10 up-regulated (right panel) Gene Ontology Biological Processes, along with Hallmark (H) pathways, with an absolute NES > 2.5. Bubble colors represent common biological processes: red for angiogenesis; green for immune pathways; orange for mitochondria processes; gray for other. p<0.05 *; p<0.0001 ****.

### Molecular and clinical characteristics of HIF-active pRCC tumors

To investigate the molecular characteristics of pRCC subgroups, we performed differential gene expression analyses of HIF-active versus Cluster A. We identified 3,132 significantly upregulated and 2,375 downregulated genes (Supplementary Table 4). Gene set enrichment analysis revealed that HIF-active tumors overexpressed genes in immune response pathways, vascular processes, and cell cycle/proliferation and had lower expression of genes involved in mitochondrial functions, particularly related to Krebs cycle and oxidative phosphorylation (Fig. 1D, Supplementary Table 5).

Transcriptomic classification of pRCC tumors according to their TME profiles, revealed that HIF-active tumors were enriched in angiogenesis-related signatures, similar to FH/SDHB-deficient tumors, while Cluster A tumors had significantly lower angiogenesis scores (P<0.0001; Fig. 2A-2B). Indeed, the vascular endothelial growth factor A (VEGFA), a well-established HIF target, is highly expressed in both HIF-active pRCC and FH/SDH-deficient tumors, reinforcing the evidence of HIF activation. Regarding immune cell infiltration, RNA-seq data deconvolution revealed that the HIF-active group is characterized by the highest immune cell infiltration, surpassing the pseudohypoxic control group (Fig. 2D). CD8 T lymphocytes, regulatory T cells, and naïve B cells, were among the abundant immune cells with the highest differences between clusters (absolute scores > 0.03 and fold change > 2.25; Fig. 2E, Fig. S5A-B).

**Figure 2.**
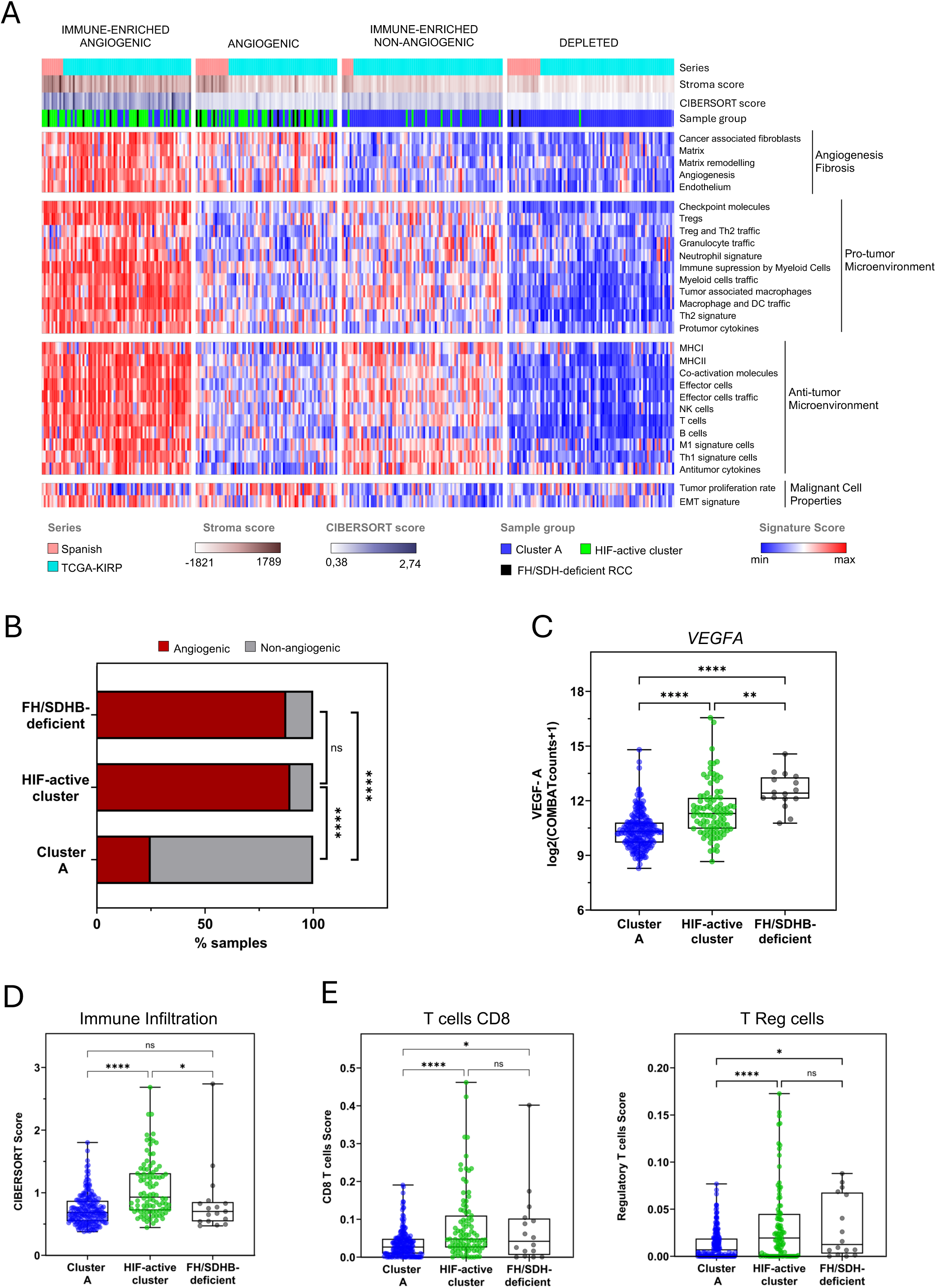
HIF-active tumors exhibit an angiogenic and immune-infiltrated tumor microenvironment. **A)** Classification of tumors into the four distinct TME subtypes described by Bagaev et al. 2021, based on unsupervised k-means clustering of the 29 Fge enrichment scores. Annotation of Series, FH/SDHB status, HIF cluster, Stroma Score and CIBERSORT are exhibited as depicted in the legend **B)** Bar chart representing angiogenic TME. Two-sided Fisher’s exact test. **C)** mRNA levels of *VEGFA* gene. Tukey’s multiple comparisons test. **D)** Scatter plot of CIBERSORT Absolute Scores. Two-sided T test. **E)** Scatter plot of the two most abundant immune cell populations: T cells CD8 (left panel) and Regulatory T cells (right panel). Two-sided T-test. p>0.05, ns; p<0.05, *; p<0,01, **; p<0.0001, ****.

Regarding clinical characteristics, HIF-active pRCC cases exhibited higher aggressivity than those in cluster A, with more patients diagnosed at stages III-IV (42% versus 19%, for HIF-active and cluster A, P<0.0001, respectively; Fig. 3A), and more frequent metastatic disease (32% versus 18%, for HIF-active and Cluster A, respectively, P=0.02; Fig. 3B). This agrees with the overexpression of the EMT pathway and elevated EMT scores detected in HIF-active tumors (Fig. S5C-D). Furthermore, HIF-active patients had shorter overall survival (P<0.001, Fig. 3B). This association remained significant in multivariate analyses correcting for multiple potential confounders, as such, age, sex and stage at diagnosis (Table 2). These findings underscore the aggressive nature of HIF-active pRCC.

**Figure 3.**
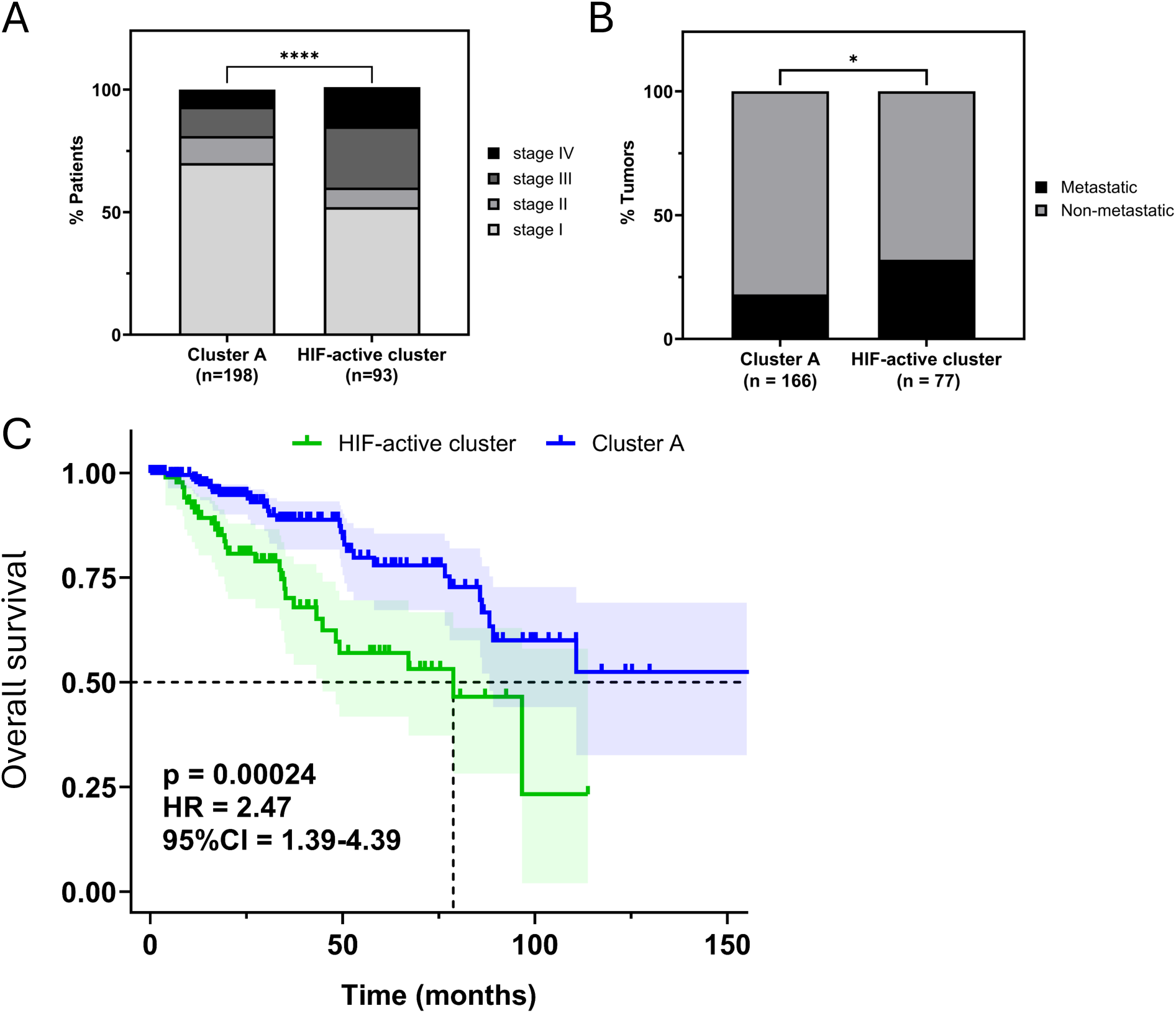
HIF-active pRCC tumors are associated with higher aggressiveness and worse OS. **A)** Bar chart representing the percentage of patients diagnosed at stages I, II, III and IV in each group. Chi-square test. **B)** Bar chart representing the percentage of patients who developed metastasis in each group. Two-sided Fisher’s exact test. **C)** Kaplan-Meier curves showing patient overall survival according to the pRCC cluster. Logrank test. 3 patients remain in Cluster A after 150 months. p<0.05, *; p<0.0001, ****. HR=Hazard Ratio, CI=Confidence Intervals.

**Table 2.** Cox regression analysis of overall survival.

| **Characterists** | **HR (95% CI)** | **p value** |
| --- | --- | --- |
| **Univariate** |  |  |
|  | 2.69 (1.59 – 4.57) | <0.001 |
| **Multivariate<sup>a</sup>** |  |  |
| HIF-active cluster | 2.81 (1.64 – 4.84) | <0.001 |
| Age | 1.02 (0.99 – 1.04) | 0.185 |
| Sex | 1.34 (0.72 – 2.69) | 0.384 |
| HIF-active cluster | 1.98 (1.13 – 3.51) | 0.018 |
| Stage<sup>b</sup> | 2.32 (1.82 – 2.99) | <0.0001 |
| Age | 1.02 (0.99 – 1.05) | 0.199 |
| Sex | 1.85 (0.98 – 3.77) | 0.070 |
| HIF-active cluster | 1.98 (1.01 – 3.57) | <0.05 |
| Stage II | 2.38 (0.84 – 5.90) | 0.076 |
| Stage III | 2.65 (1.23 – 5.58) | <0.05 |
| Stage IV | 15.47 (7.68 – 31.88) | <0.0001 |
| Age | 1.02 (0.99 – 1.05) | 0.174 |
| Sex | 1.82 (0.95 – 3.74) | 0.084 |

HR = Hazard Ratio; CI = Confidence interval
<sup>a</sup>Multivariate analysis including together with the pRCC cluster different clinical variables (age, sex, stage at diagnosis).
<sup>a</sup>Stage at diagnosis included in the analysis as an ordinal variable from I to IV.

### L-2-Hydroxyglutarate dehydrogenase downregulation drives HIF activity

Next, we sought to explore the mechanisms behind HIF overactivation in pRCC. Fumarate, succinate and L-2-hydroxyglutarate (L-2-HG) are oncometabolites that drive RCC tumorigenesis through the inhibition of α-ketoglutarate-dependent dioxygenases^42^. The accumulation of these oncometabolites in the tumors leads to HIFα accumulation through HIF PHDs inhibition and could explain pRCC HIF-active cluster characteristics. In addition, fumarate and succinate lead to global DNA hypermethylation (CIMP) through TET enzyme inhibition^42,43^. To investigate if these oncometabolites could play a role in HIF-active tumors, we examined the methylome data of the samples. While FH-mutated tumors showed strong DNA hypermethylation, HIF-active and Cluster A tumors did not. Although the HIF-active showed higher methylation levels than Cluster A (Fig. 4A-B). These results rule out fumarate and succinate accumulation as causative of HIF activation in pRCC. Regarding L-2-HG, the inhibition capacity of this metabolite on TET enzymes is 10-fold lower than fumarate and succinate, suggesting that cellular concentrations of L-2-HG are insufficient to cause CIMP^42^, but could explain the differences observed among the pRCC clusters.

**Figure 4.**
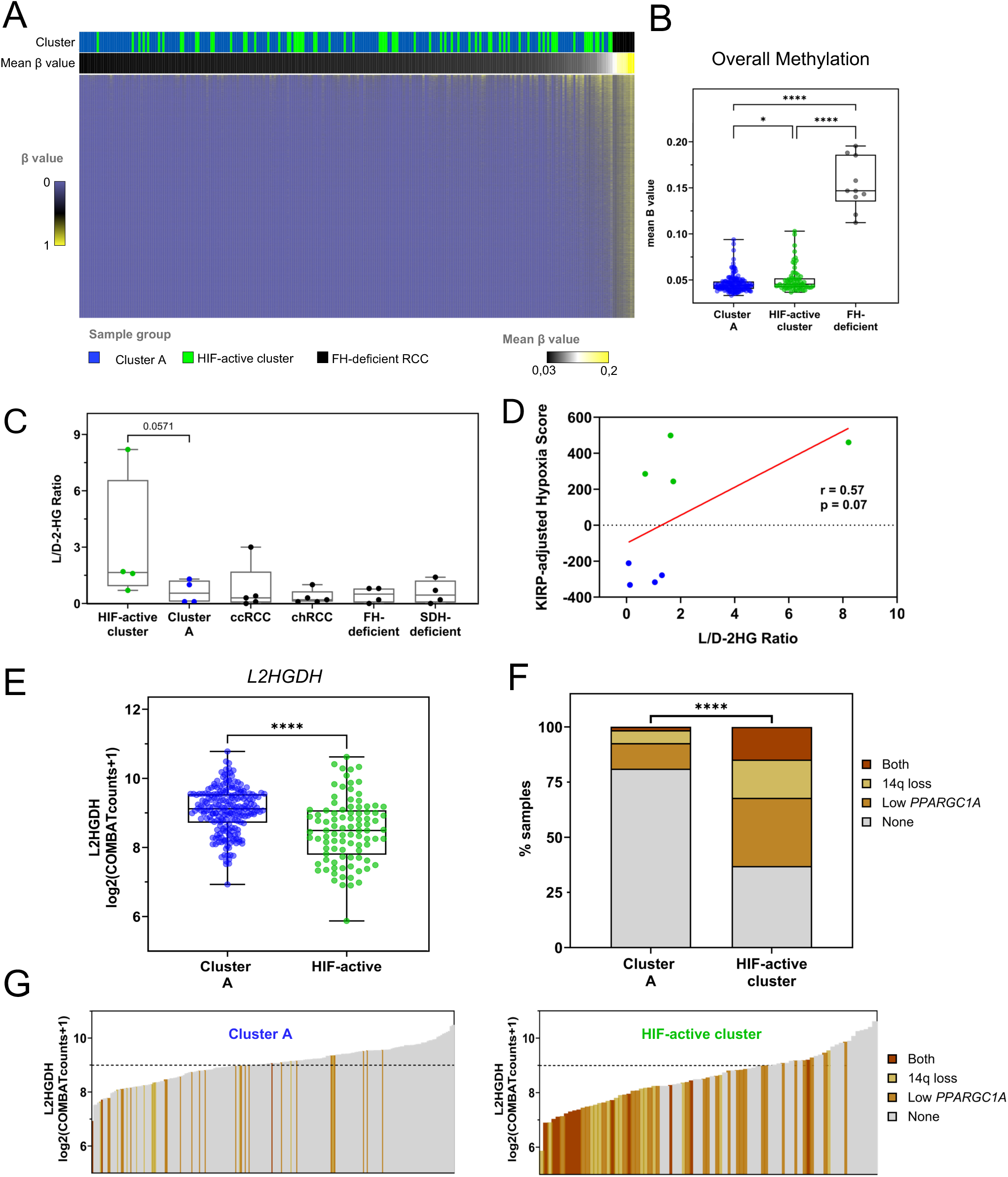
Methylation level, L-2-HG accumulation and mechanisms of *L2HGDH* downregulation in HIF active tumors. **A)** Heatmap representation of the DNA methylation level of 14,915 unmethylated probes in normal tissue. Tumors were annotated for sample group and sorted by mean methylation level (β value). **B)** Scatter plot of mean β values. Tukey’s multiple comparisons test. **C)** L/D-2-HG ratios in RCCs. One-tailed Mann Whitney test. **D)** Correlation between L/D-2-HG ratio and KIRP-adjusted hypoxia score. One-tailed Pearson correlation. **E)** *L2HGDH* expression levels. Two-sided T test. **F)** Bar chart representing the percentage of samples with each alteration. Samples belonging to the first quartile *PPARGC1A* expression are classified as *Low PPARGC1A.* Chi-square test. **G)** Waterfall plots depicting the distribution of *L2HGDH* expression according to each alteration in Cluster A (left panel) and in HIF-active cluster (right panel). The dashed line indicates the median *L2HGDH* expression in samples. p<0.05, *; p<0.0001, ****.

To determine if L-2-HG accumulation could be the cause of HIF overactivation in HIF-active cluster, we performed a metabolomic analysis in an RCC series that included pRCC cases from our Spanish cohort and other RCC histologic subtypes. As shown in Fig. 4C, the highest L-2-HG/D-2-HG ratio corresponded to HIF-active tumors, with a 2.9-fold increase versus Cluster A and a positive correlation with the hypoxia score of the pRCC (r=0.57; Fig. 4D). As expected, fumarate and succinate were accumulated in FH and SDH-deficient RCC, and interestingly, the glutamate/aspartate ratio (associated with HIF activity^44,45^) was maximum in ccRCC, followed by FH-RCC, HIF-active pRCC, and SDH-RCC (Fig. S6A-B).

As a downregulation of L-2-hydroxyglutarate dehydrogenase (*L2HGDH*) expression is responsible for L-2-HG accumulation in cancer^46,47,48^ to further confirm the causative role of L-2-HG in HIF-active tumors, we determined the expression of this gene in the pRCC series. As shown in Fig. 4E, indeed, *L2HGDH* expression was significantly lower in HIF-active versus Cluster A (P<0.00001). Furthermore, *L2HGDH* expression had an inverse correlation with the L/D-2-HG ratio and with the pRCC hypoxia score (Fig. S7A-B).

The alterations leading to a low expression of *L2HGDH* in RCC include a loss of the 14q chromosome arm^46,47,48^ where the gene is located, and a reduced expression of the transcription factor PGC1α^48^, encoded by the *PPARGC1A* gene. We observed that these were common events in pRCC, and both alterations, individually or together, were more frequent in the HIF-active group than in Cluster A (63% of HIF-active tumors presented one or both alterations, compared to 19% in Cluster A; P<0.00001; Fig. 4F). *PPARGC1A* expression had a positive correlation with *L2HGDH* and was lower in HIF-active group (Fig. S7C-D). As shown in Fig. 4G and Fig. S7E, these alterations in pRCC were associated with reduced expression of *L2HGDH* and the increased pseudohypoxic phenotype.

## DISCUSSION

The extensive molecular heterogeneity of pRCC could underlie the large variability in response to standard targeted therapies^1,2,6–9^, emphasizing tumor stratification as a critical feature to advance precision medicine in this disease. In this study, by analyzing a large cohort of 303 pRCC tumors enriched in metastatic cases, we identified for the first time a subgroup of papillary tumors exhibiting elevated HIF activity. These tumors have a unique tumor microenvironment and poor clinical outcomes. We propose the accumulation of L-2-HG, an unexplored mechanism in pRCC, as the primary cause driving HIF activity (Fig. 5).

**Figure 5.**
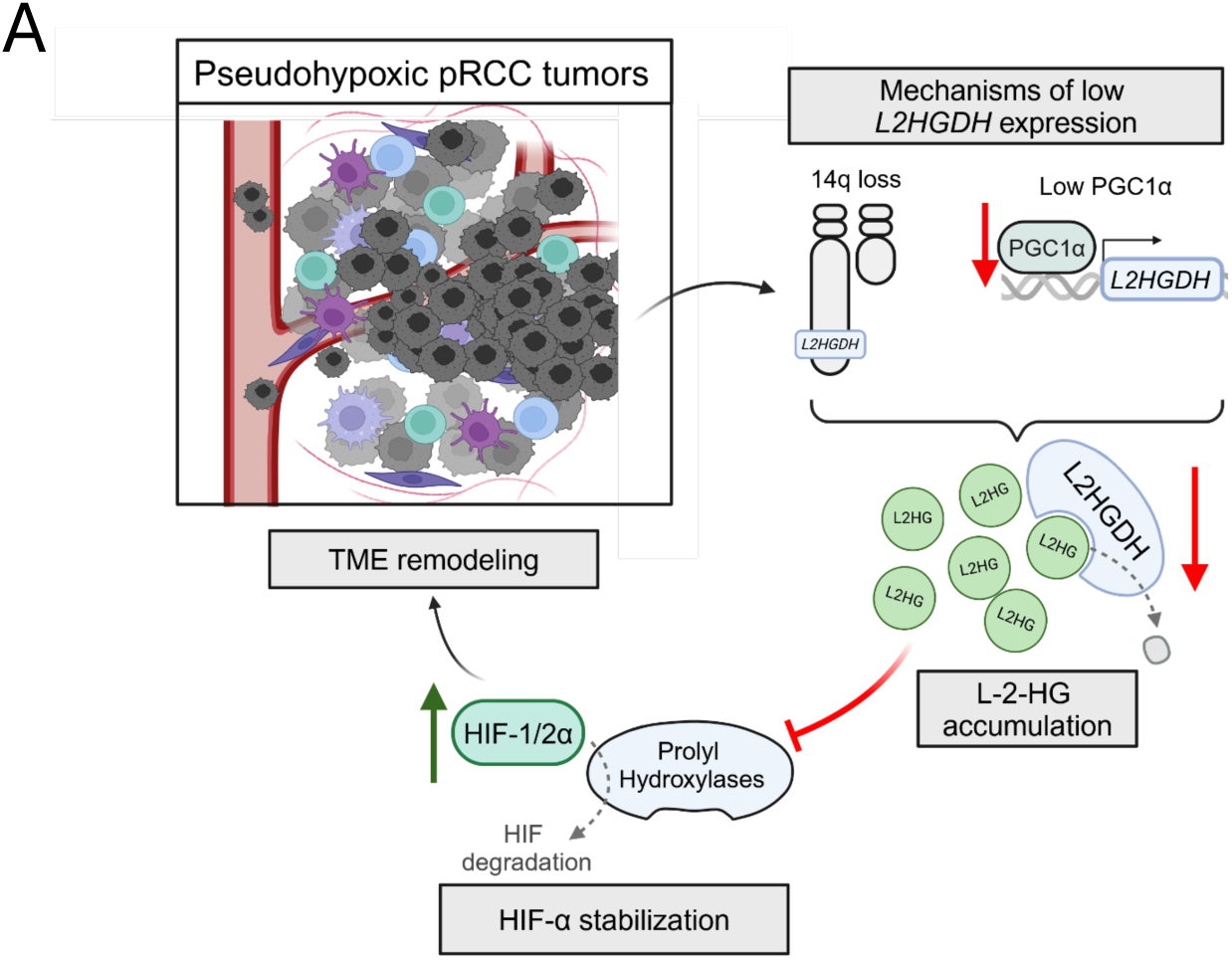
Graphical abstract illustrating the downregulation of *L2HGDH* and its subsequent effects.

The critical role that HIF plays in tumorigenesis and tumor microenvironment reshaping has been widely described across various cancers^14,15,49–52^, particularly in ccRCC^4,53^. In agreement, the HIF-active pRCC subgroup identified here showed a high hypoxia score, a shift towards non-oxidative metabolism, upregulation of proliferative pathways and increased EMT. Changes in the tumor microenvironment included increased angiogenesis and immune cell infiltration. These characteristics align with an over-activated HIF pathway, which promotes the expression of genes that enhance oxygen delivery, reduce oxygen consumption, and support cell survival under hypoxic conditions^10–13,52^. The high tumor immune infiltration may also be attributed to HIF signaling, which promotes not only vasculature development but also lymphangiogenesis^52,54^. HIF-active pRCC are immune “hot” tumors, suggesting sensitivity to immune checkpoint inhibitors. High quantities of cytotoxic cells, such as NK and CD8+ T cells, along with macrophages, can promote an anti-tumor microenvironment, however, the presence of regulatory T cells and anti-inflammatory M2 macrophages, alongside a high proportion of NK and dendritic cells in a resting state can lead to a general immunosuppressive environment (Fig. S4A-B). Thus, further investigation is needed to fully understand the immune phenotype of HIF-active pRCC.

Mechanisms that explain HIF activation in tumors under normoxic conditions (i.e. pseudohypoxia) include gain-of-function mutations in *EPAS1* gene^55^, loss of pVHL^10,11,25,67^, and the accumulation of various oncometabolites due to mutations in the Krebs cycle genes (*FH*, the *SDH* genes and *IDH*)^15,42^. Furthermore, while *L2HGDH* mutations are rare in cancer, a downregulation in the expression of this enzyme lead to the accumulation of L-2-HG^46–48^, another oncometabolite capable of inhibiting several α-ketoglutarate dependent dioxygenases. These dioxygenases include not only HIF-prolyl hydroxylases, but also HIF Factor Inhibiting 1, another crucial regulator of HIF degradation^42,56^. Regarding TET1/2 demethylases, these enzymes are heavily inhibited by fumarate, succinate and D-2-HG, resulting in the so-called CIMP^42,43^. However, whether the accumulated amounts of L-2-HG observed in tumors can inhibit TET1/2 is unclear^42,46^, in line with the small increment in methylation observed in our pRCC HIF-active tumors.

This study is the first to discover that *L2HGDH* expression is low in a subset of pRCC, which corresponds to HIF-active cluster. In agreement, a metabolomic analysis showed a high L-2HG/D-2HG ratio in HIF-active tumors compared to other types of RCCs. The mechanisms causing decreased *L2HGDH* expression in ccRCC (loss of chromosome 14q and/or downregulation of *PPARGC1A*) were also enriched in the HIF-active pRCC subset.

Chromosome 14q also harbors *HIF1A* gene and may suggest shifting HIF activity towards HIF-2α dependency and sensitivity to HIF-2α inhibitors. While HIF-1α is widely recognized as a central regulator of glycolytic metabolism, angiogenesis, and survival pathways, helping tumors adapt to hypoxic conditions, HIF-2α exhibits a more tissue-specific role, contributing to the maintenance of stemness and promotion of EMT, thereby facilitating metastasis. Interestingly, the prognostic value of these factors is still contradictory, as their impact on outcomes varies largely across cancer types^57^. In our pRCC series, although HIFα are regulated mainly at the protein level, transcriptional data show greater differences between HIF-active and Cluster A for *EPAS1* than for *HIF1A* expression (encoding HIF-2α and HIF-1α, respectively). This would support the hypothesis of tumor dependence on HIF-2α, whose transcription would be enhanced to maintain high levels of this factor. Therefore, future analysis of HIF protein expression could help distinguish which HIF isoform is the main driver of the HIF-active pRCC.

The subset of HIF-active pRCC cases is 31% in this study, but considering their aggressive behavior, this figure could be higher among metastatic pRCC. In fact, if we consider only the metastatic cases in our series, this figure increases to 45%. HIF active tumors exhibit an inflammatory and angiogenic tumor microenvironment, suggesting that these cases may be the ones more likely to respond to current recommended first-line treatments for metastatic pRCC (VEGFR-TKI monotherapy, immunotherapy, or a combination of both)^6–9^. In addition, the multiple effects triggered by L-2-HG accumulation, including its impact on anti-tumor immunity^58^ and metabolic dysregulation, represent a new opportunity to explore innovative and specific therapeutic strategies in this cancer subtype.

Despite the limitations of the study in terms of sample size and series heterogeneity, challenges inherent to research on low-frequency diseases, we performed a robust histologic subtype analysis to adhere to the new WHO 2022 RCC tumor classification and made a significant effort to molecularly dissect pRCC tumors and identify distinct characteristics and prognostic implications.

The need for molecular stratification in pRCC is becoming increasingly evident, with putative clinical impact highlighted by this study. This newly identified HIF-active tumor subgroup and its unique microenvironment features may account for the variable effectiveness of standard RCC therapies, and supports the rationale for combining immunotherapy with anti-angiogenic drugs in this specific group of patients, thereby opening a window for more precise treatment approaches in metastatic pRCC.

## ACKNOWLEDGEMENTS

We thank CNIO Histopathology Unit for their support in FFPE tumor tissue processing and hematoxylin-eosin staining, Rocío Letón for her help in genetic techniques and Angel Martinez-Montes, Solip Park and Manuel Moradiellos for their bioinformatic support.

This work was supported by the grants PID2021-128312OB-I00, funded by MCIN/AEI/10.13039/501100011033 (CRA) and the Spanish Ministry of Science, Innovation and Universities “Formación del Profesorado Universitario – FPU” fellowships with ID numbers FPU21/02012 (JNH), and FPU20/02798 (CV).

## DISCLOSURE OF CONFLICT OF INTEREST

None.

**Figure S1.**
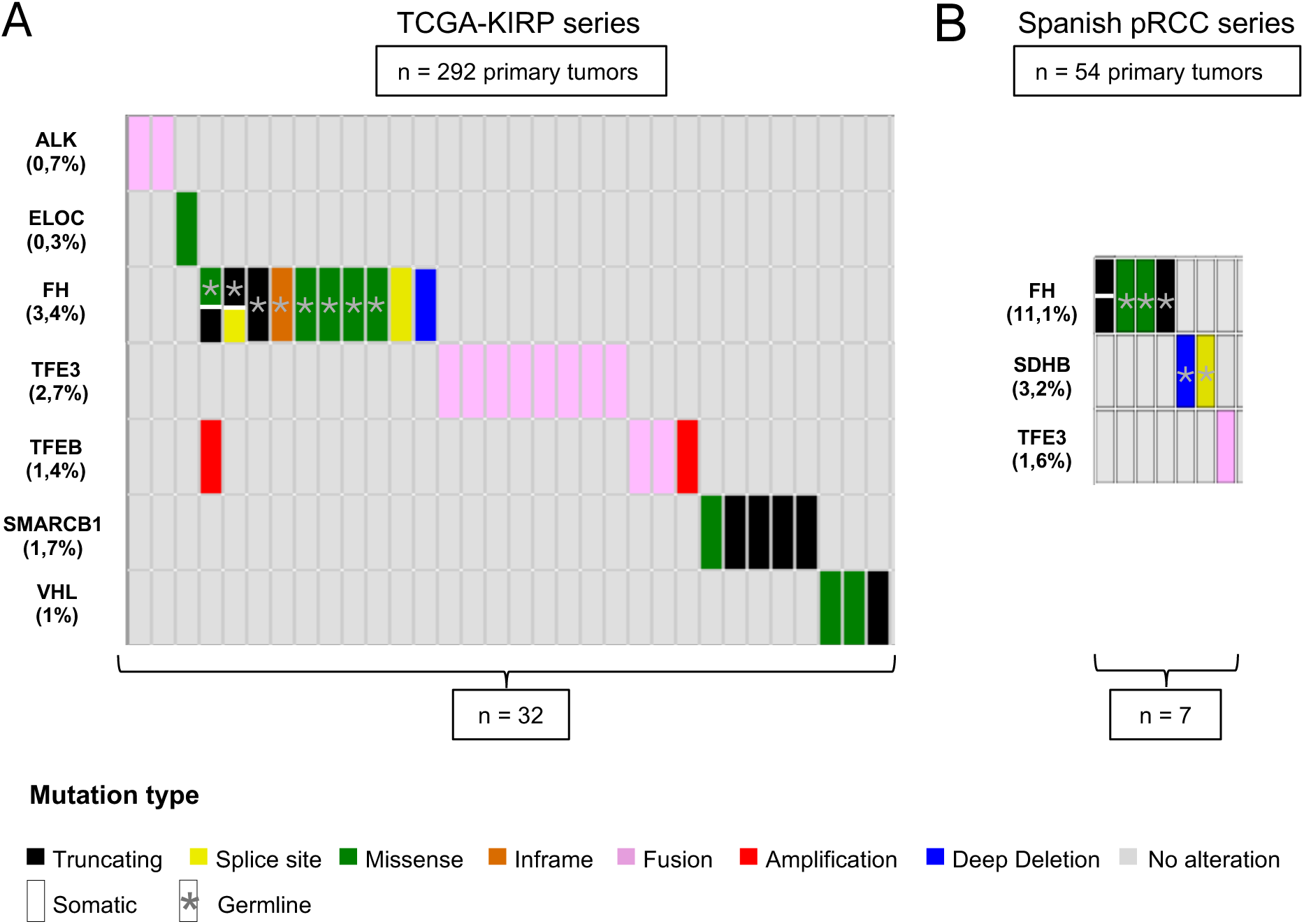
Somatic and germline driver mutations identified and used for RCC histology reclassification. Oncoprint of driver alterations in **A)** TCGA-KIRP series and **B)** Spanish pRCC series. Cell division means a sample with two mutations. Grey asterisks denote germline mutations.

**Figure S2.**
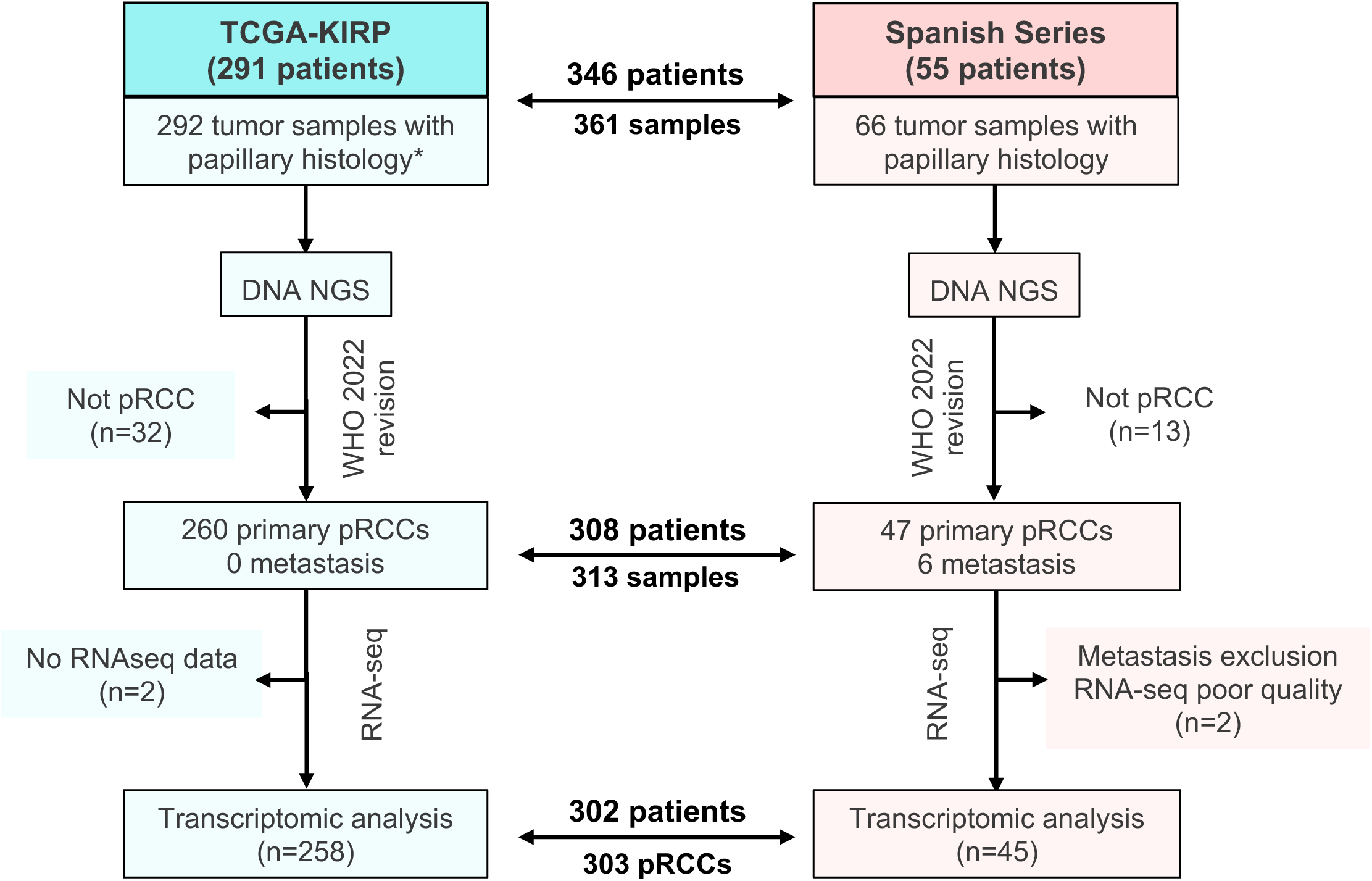
Flow chart showing the final number of independent pRCC samples included in the transcriptomic analysis. The diagram presents the number of independent tumor samples (defined as those obtained in different surgeries) in the Spanish series and the TCGA-KIRP included in the final transcriptomic analysis. For pRCC classification, samples had to pass a mutational revision based on DNA NGS. To be included in the final transcriptomic analysis, samples had to pass RNAseq quality control. *One patient had two independent primary tumors

**Figure S3.**
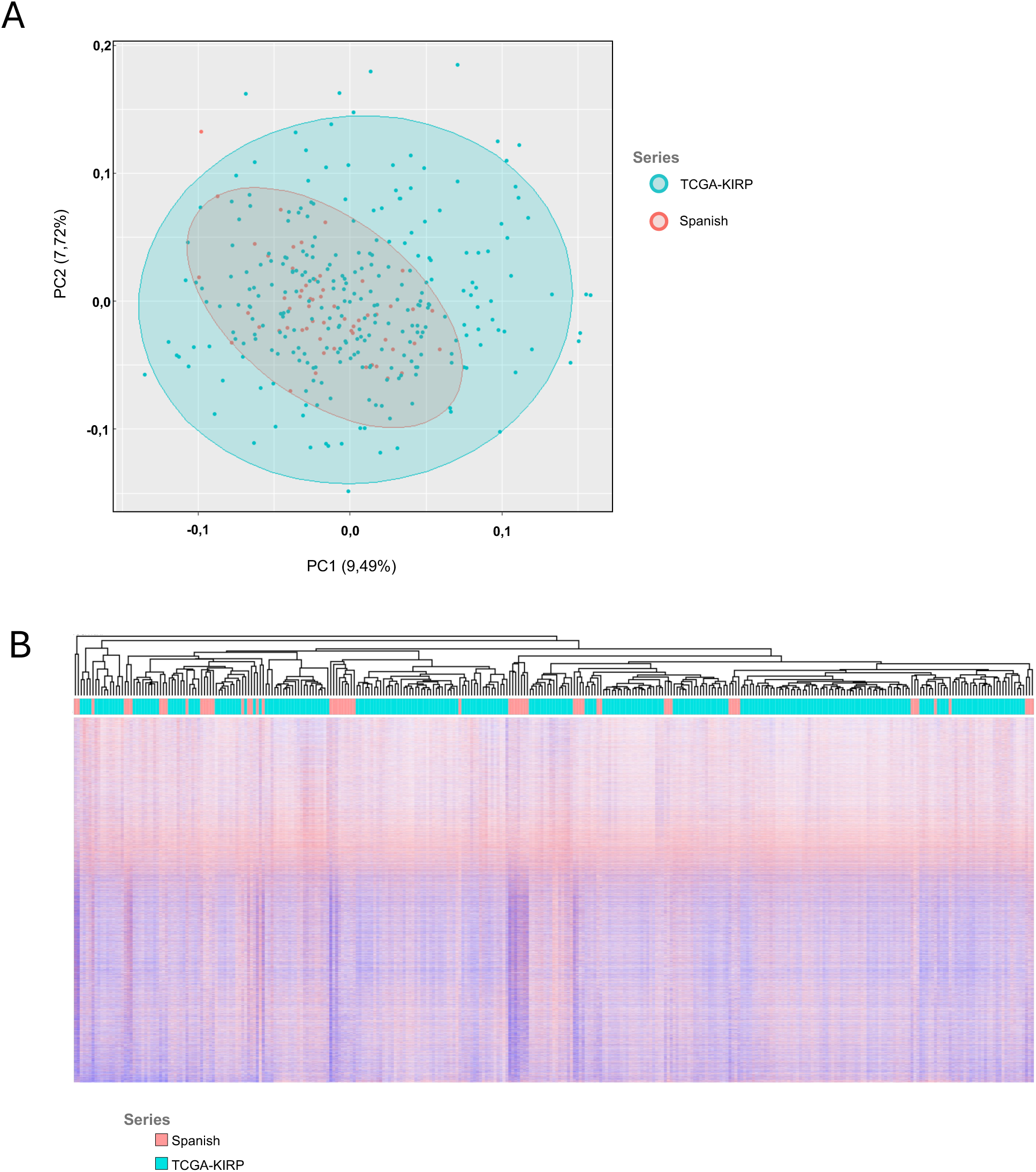
Integration of RNA-seq data of 303 pRCC tumors from both TCGA-KIRP and Spanish series. **A)** Principal Component Analysis based on the expression of 22,835 genes of tumors after COMBAT-seq correction. **B)** Heatmap of unsupervised hierarchichal clustering using the expression of 22,835 genes after COMBAT-seq correction.

**Figure S4.**
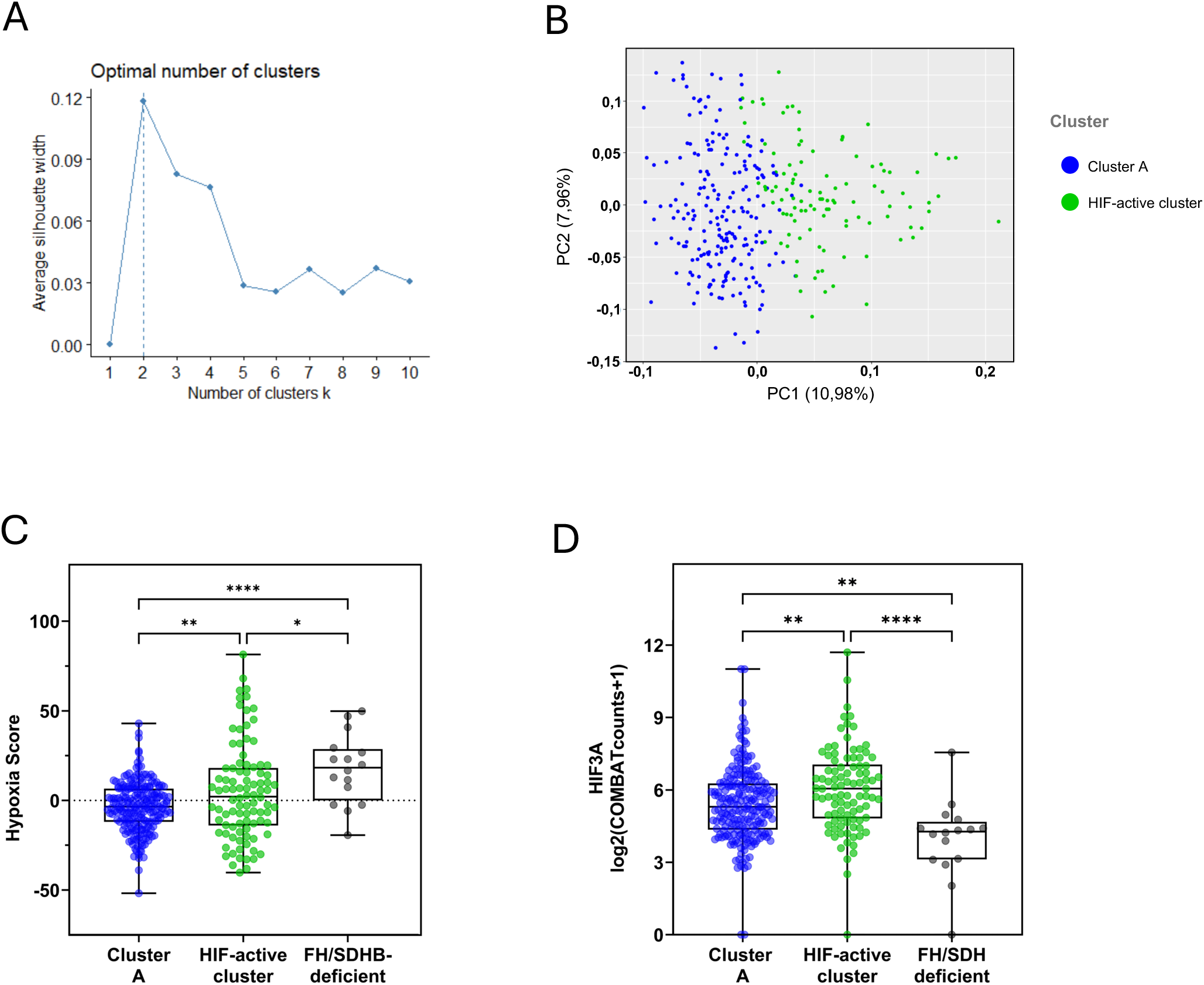
Combination of RNA-seq data and some features of HIF target genes-based clusters. **A)** Optimal number of clusters (K=2) determined by the Silhouette method based on the expression of 461 HIF target genes. **B)** Principal Component Analysis based on the expression of the HIF-targets, colored by cluster assignation (HIF-active cluster, light green; Cluster A, dark blue). **C)** Hypoxia Score calculated as described by Lombardi et al. 2022 (PMID: 36384128). **D)** mRNA levels of *HIF3A* gene. Tukey’s multiple comparisons test. p<0.05, *; p<0,01, **; p<0.0001, ****.

**Figure S5.**
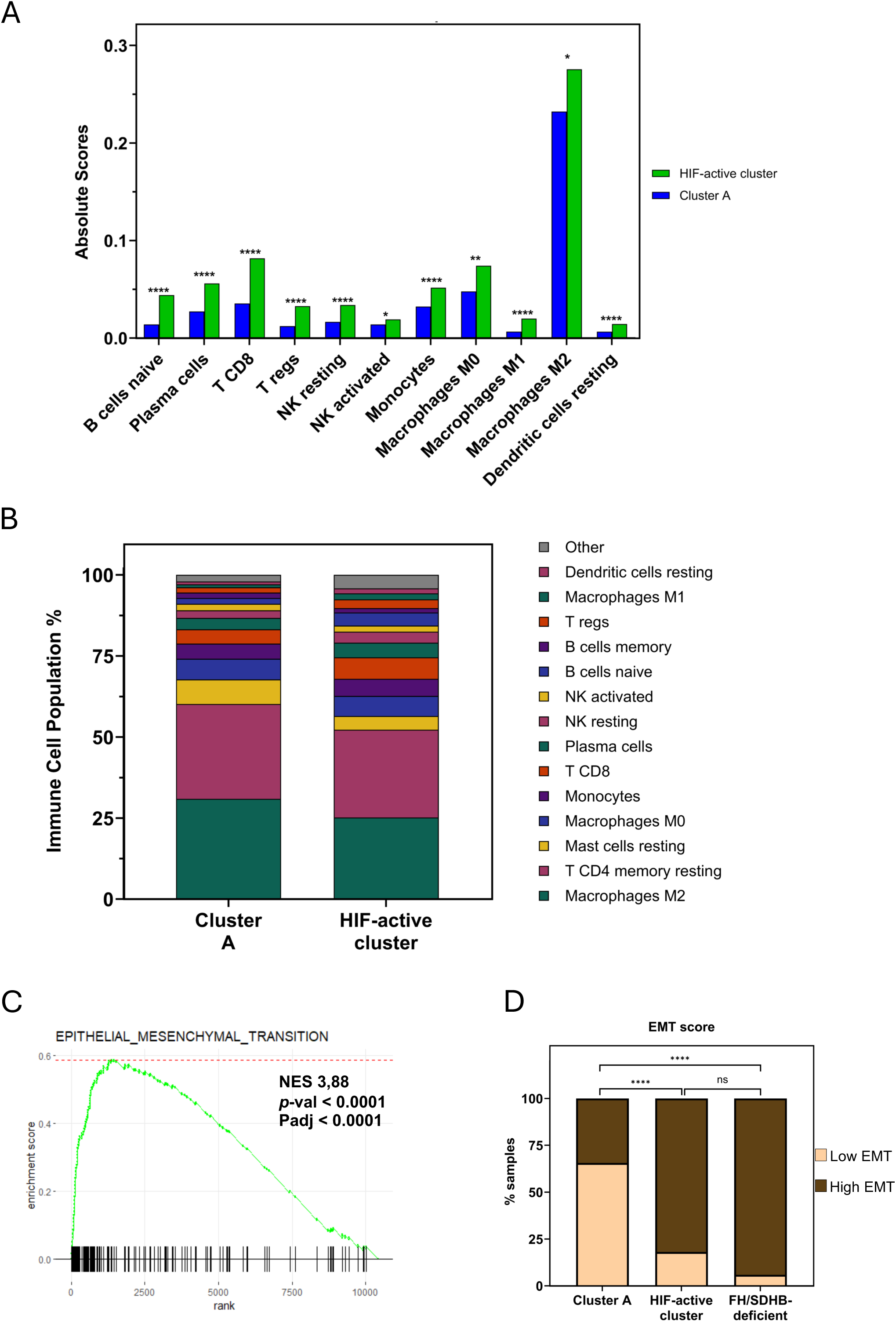
HIF-active tumors exhibit high immune cell infiltration and metastatic signatures. **A)** Mean absolute scores of immune cell populations derived from CIBERSORTx deconvolution. Two-tailed t-test. **B)** Bar plot depicting the percentage of most abundant immune cell populations. **C)** GSEA plot of Hallmark Epithelial to Mesenchymal Transition (EMT) gene set. **D)** Bar plot comparing percentage of samples with high and low EMT score, calculated as described in Aggarwal RK et al., 2021. Tukey’s multiple comparisons test. p>0.05, ns; p<0.05, *; p<0,01, **; p<0.001, ***; p<0.0001, ****. NES = Normalized Enrichment Score. p-val = p-value. Padj = adjusted p-value.

**Figure S6.**
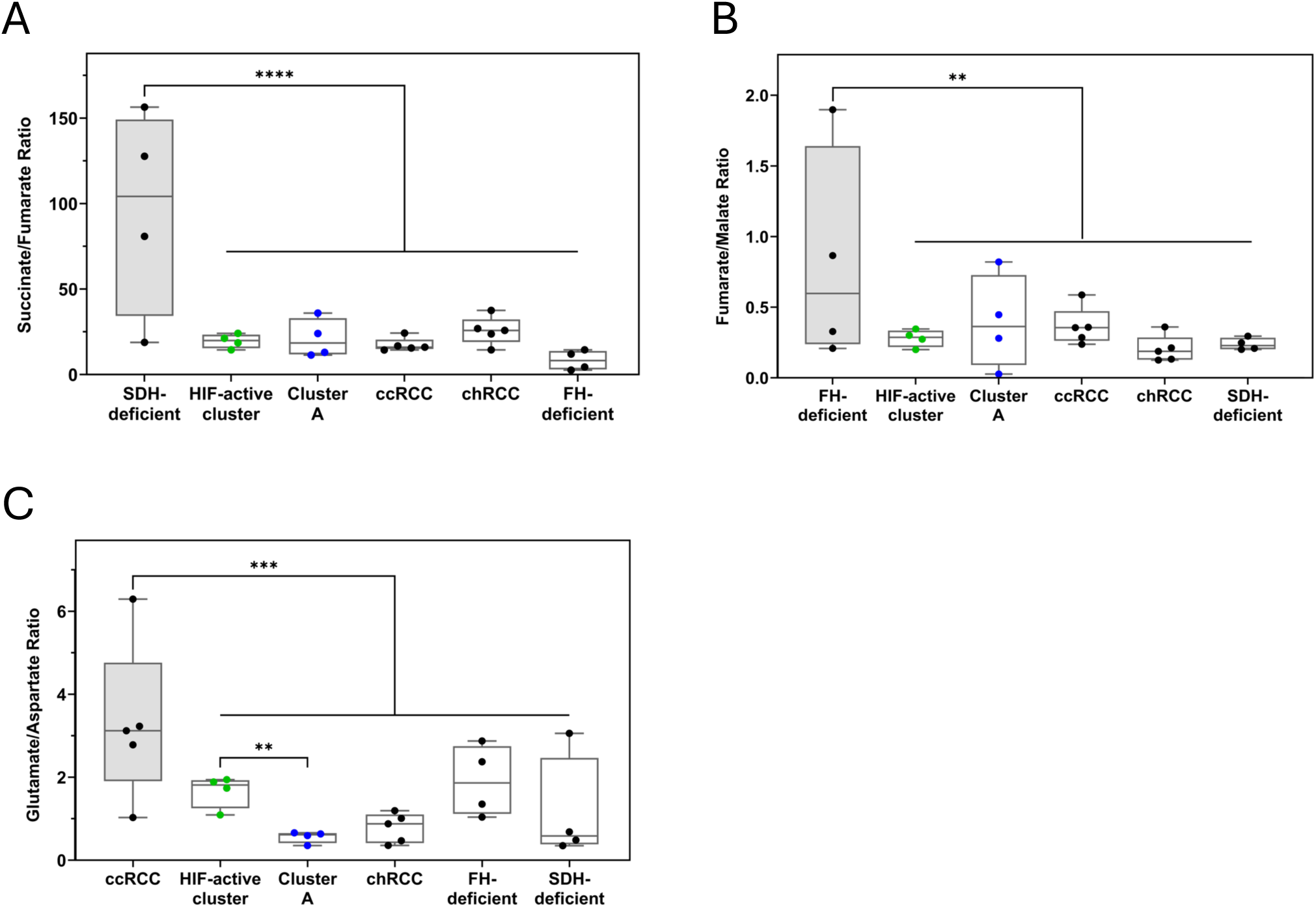
Metabolic analyses reveal alterations in glutamate and aspartate levels, but no effect in fumarate or succinate levels. **A)** Succinate/Fumarate ratios of analyzed RCCs. Dunnett’s multiple comparisons test. **B)** Fumarate/Malate ratios of analyzed RCCs. One-tailed Mann Whitney test **C)** Glutamate/Aspartate ratios of analyzed RCCs. One-tailed Mann Whitney test. p<0.01, **; p<0.001, ***; p<0.0001, ****. Reference group for each ratio is located on the left of the graph and colored in gray.

**Figure S7.**
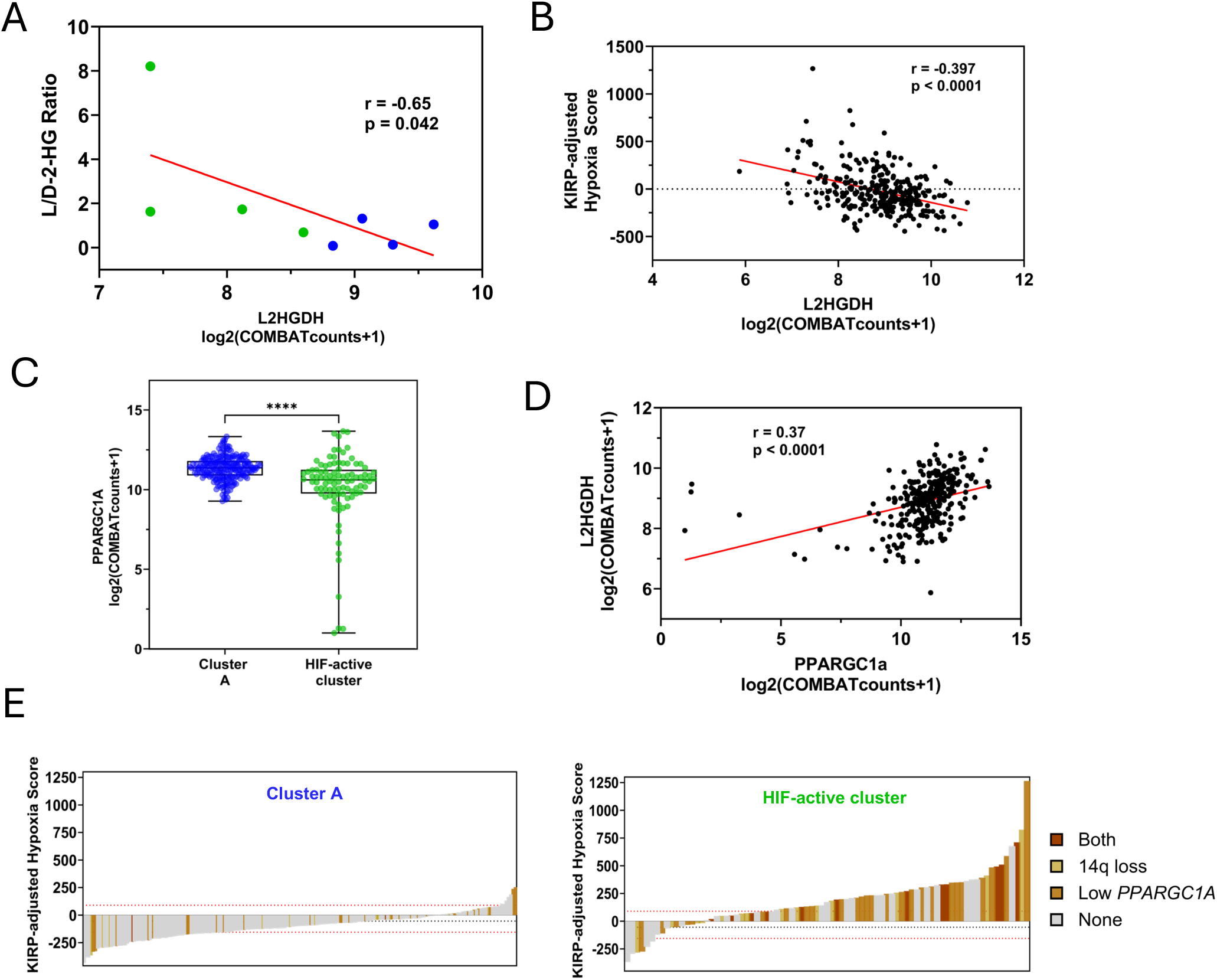
Specific molecular features of HIF-active and Cluster A tumors related to *L2HGDH* downregulation. **A)** Correlation betwewen *L2HGDH* expression and L/D-2-HG ratio. One-tailed Pearson correlation **B)** mRNA levels of *PPARGC1A* in HIF-active tumors and Cluster A. Two-sided T test. **C)** Correlation betwewen *L2HGDH* expression and the KIRP-adjusted Hypoxia Score. Two-tailed Pearson correlation. **D)** Correlation between *PPARGC1A* and *L2HGDH* expression. Two-tailed Pearson correlation. **E)** Waterfall plots depicting the distribution of Hypoxia Score according to each mechanism in Cluster A (left panel) and HIF-active cluster (right panel). Black dashed line indicates median hypoxia Score, whereas red dashed lines correspond to first and third quartiles. p<0.0001, ****.

